# Annotation and visualisation of parasite, fungi and arthropod genomes with Companion

**DOI:** 10.1101/2024.02.19.580948

**Authors:** William Haese-Hill, Kathryn Crouch, Thomas D. Otto

**Affiliations:** School of Infection & Immunity, University of Glasgow, UK; LPHI, CNRS, INSERM, Université de Montpellier, France

## Abstract

Although sequencing genomes has become increasingly popular, there is still a bottleneck for the annotation of the resulting assemblies. Structural and functional annotation is still challenging as it includes finding the correct gene sequences, annotating other elements such as RNA and being able to submit those data to databases to share it with the community. We developed the Companion web server to allow non-experts to annotate their genome using a reference-based method, enabling them to analyse their results before submitting to public databases. In this update paper, we describe how we included novel methods for gene finding and made the server more efficient to annotate genomes of up to 1 GB in size. The reference set was increased to genomes from the fungi and arthropod kingdoms. We show that Companion outperforms existing comparable tools.

**GRAPHICAL ABSTRACT:** 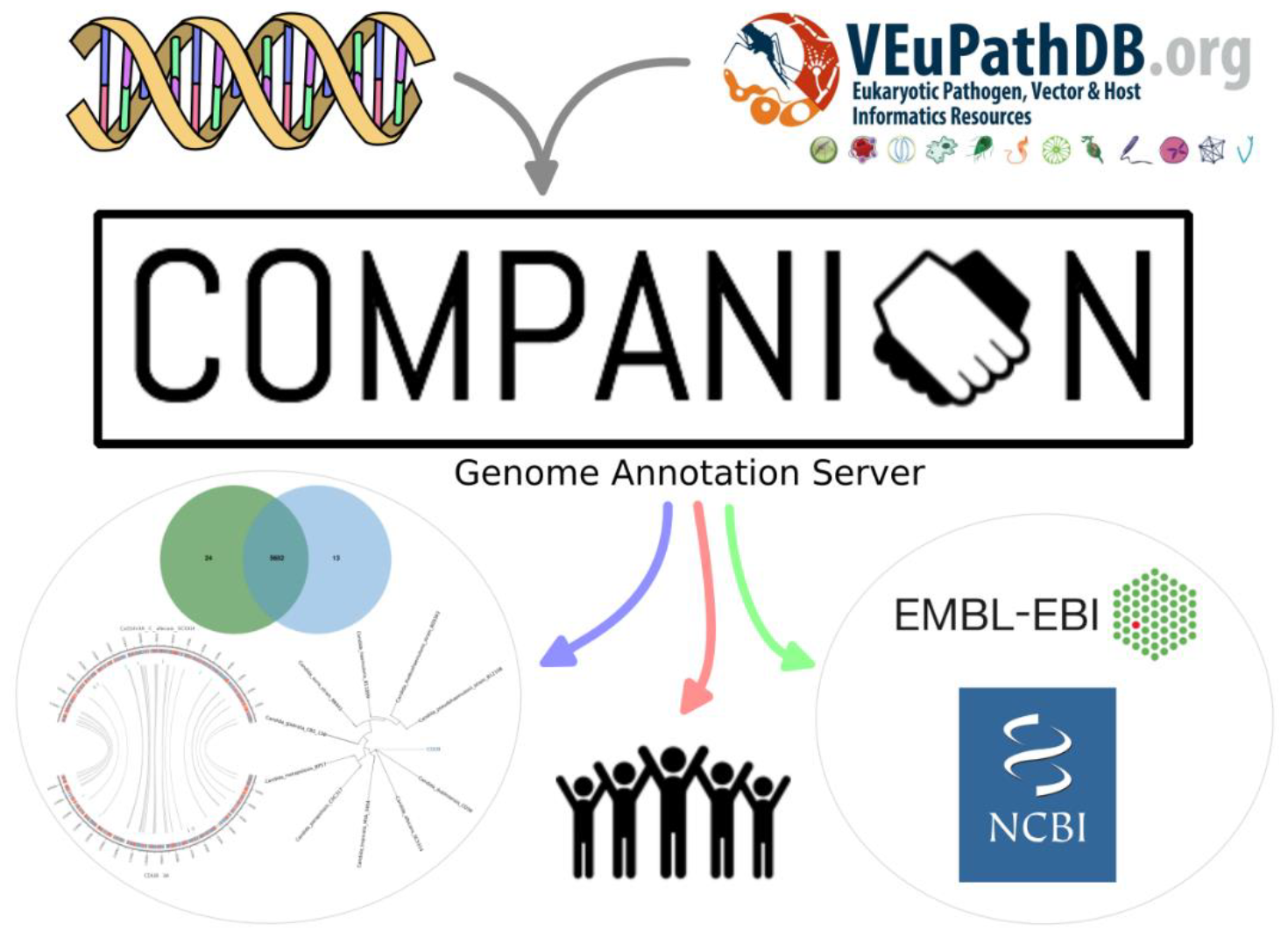

## INTRODUCTION

Over the last 15 years, the maturation of long-read sequencing technologies, the decreasing cost of sequencing, together with the development of simple and efficient software for de novo assembly (Chin, et al., 2013; Koren, et al., 2017) has enabled the community to produce continuous assemblies for species with repetitive or low complexity genomes that were fragmented and incomplete with short-read technologies. These advances inspired ambitious proposals such as the Earth BioGenome Project (EBGP) to sequence ∼1.5 million of the current estimated 10-15 million Eukaryotic species (Lewin, et al., 2018).

Although it has become increasingly easy to assemble genomes, the annotation of assemblies remains a difficult problem and is often neglected. For example, the exon accuracy of common annotation tools used to determine gene models, such as MAKER2, ranges only between 55-70% (Table 1 in Holt and Yandell, 2011). This difficulty arises due to the complexity of the biological system, where genes can be structured differently depending on the organism. Genes have alternative splicing and even with the use of long read transcriptomics sequencing does not result in high quality annotations. Further, in a full annotation process, many different tools are involved; whether gene finding, functional annotation or ncRNA detection. There are additional challenges in submitting the annotations to International Nucleotide Sequence Database Collaboration (INSDC) (GenBank, EBI and DDBJ) due to an overly complex pipeline for uploading the data. This results in poor public availability of genome annotations and data that does not uphold Findable, Accessible, Interoperable, and Reusable (FAIR) principles (Barker, et al., 2022).

**Table 1.** Standard metrics to compare performance of GenSAS and Companion against canonical annotations, for various organisms. Accuracy is mean of sensitivity and precision values. Note significant improvement in exon accuracy and matched loci for Companion in all cases, as well as greater number of total genes predicted. † comparison of CDS features only due to presence of UTR features in canonical annotation ‡ removed UTR features from Companion output due to absence in canonical annotation * times given for full pipeline and minimal pipeline jobs (see Supplementary Data). Referenced statistics are from the latter

| Organism | Metric | GenSAS | Companion |
| --- | --- | --- | --- |
| *Plasmodium falciparum*‡<br>(5,642 total genes) | Nucleotide accuracy (%) | 98.65 | 99.35 |
|  | Exon accuracy (%) | 79.35 | 95.05 |
|  | Matching loci (%) | 66.64 | 96.22 |
|  | Total genes predicted | 5,034 | 5,800 |
|  | Runtime (h) | 6 / 1* | 2.5 |
| *Candida albicans*<br>(6,459 total genes) | Nucleotide accuracy (%) | 98.55 | 98.35 |
|  | Exon accuracy (%) | 85.95 | 89.50 |
|  | Matching loci (%) | 90.85 | 95.54 |
|  | Total genes predicted | 5,822 | 6,206 |
|  | Runtime (h) | 17.5 / 1* | 1 |
| *Anopheles darlingi*†<br>(11,005 total genes) | Nucleotide accuracy (%) | 87.1 | 86.15 |
|  | Exon accuracy (%) | 66.75 | 74 |
|  | Matching loci (%) | 48.27 | 65.05 |
|  | Total genes predicted | 14,429 | 16,500 |
|  | Runtime (h) | 13.5 / 3* | 9 |

Therefore, several tools have been developed to perform automated genome annotation. However, some tools, such as CAT (Fiddes, et al., 2018) and FunAnnotate (Palmer and Stajich, 2023), lack web-based or graphical interfaces, raising the barrier to tool use. Easier to use are webservers, like GenSAS (Humann, et al., 2019), MEGANTE (Numa and Itoh, 2014) or the NCBI Eukaryotic Annotation Pipeline (Thibaud-Nissen, et al., 2016). However, MEGANTE is limited by a maximum upload file size of 10MB and lacks appropriate training models for species like Fungi. The NCBI server requires a user to send an email requesting annotation, centralising the annotation process rather than democratising it. Further, looking at the statistics it seems to be at full capacity, and when we requested the annotation of a parasite genome, this was not possible. GenSAS worked without restrictions and contains a wide range of pretrained models for structural annotation. However, it lacks some parasite reference sets, does not have a graphical output and relies on expert knowledge to set up.

In 2016 we developed “Companion” (Steinbiss, et al., 2016). This service provides an easy-to-use pipeline for reference-based annotation with pre-compiled reference genomes derived from VEuPathDB (Amos, et al., 2021). Companion uses ABACAS (Assefa, et al., 2009) to scaffold contigs against a reference, RATT (Otto, et al., 2011) to transfer annotations as described above and AUGUSTUS (Stanke, et al., 2008) for *ab initio* gene prediction where annotations cannot be transferred. Companion supports the use of RNA-Seq data to improve annotations. Output can be downloaded in a selection of formats that have been designed to facilitate upload to EBI. A particular strength of Companion is the visual output which includes basic statistics, a phylogenetic tree generated against the reference set, analysis of orthology with OrthoMCL (Li, et al., 2003) and synteny maps. This allows users to quickly assess not only the quality of the annotation but also the completeness of the assembly.

Despite its small parasite user community, it was heavily used and with the increasing number of assemblies generated in the communities of vectors and arthropods, we decided to extend the range of references to those. Early testing with these additional, often much larger, reference genomes revealed numerous pipeline components that were unable to scale efficiently, and so needed updating to ensure a satisfactory outcome.

Here we describe the implementation improvements we did to not only allow the annotation of larger genomes (included in a total pool of 438 reference genomes, Supplementary Table S1), but also make the tools more robust and faster. Finally, we show that Companion generally outperforms other annotation server tools and should be considered as a tool for the community to annotate Fungi, Protozoan parasites and Arthropods.

## MATERIAL AND METHODS

### Updated workflow

Companion version 2 is implemented as a Nextflow DSL1 pipeline and the source code can be seen at https://github.com/sii-companion/companion. GitHub Actions is used for automated testing and tag/release generation, where releases follow standard semantic versioning format. For running locally, a docker container is hosted at https://hub.docker.com/repository/docker/uofgiii/companion/ with builds synced to each code release. Instructions are available on the GitHub Wiki for building a bespoke reference dataset when running a container locally. This ensure that users are not restricted to references hosted by VEuPathDB, as they are when using the web service.

Compared to the first version v1.0.2, several new features have been implemented (see Figure 1 – red) and others improved (blue). These changes allow annotation of genomes up to 3 GB, although we observed optimal performance in genomes up to 1 GB.

**Figure 1.**
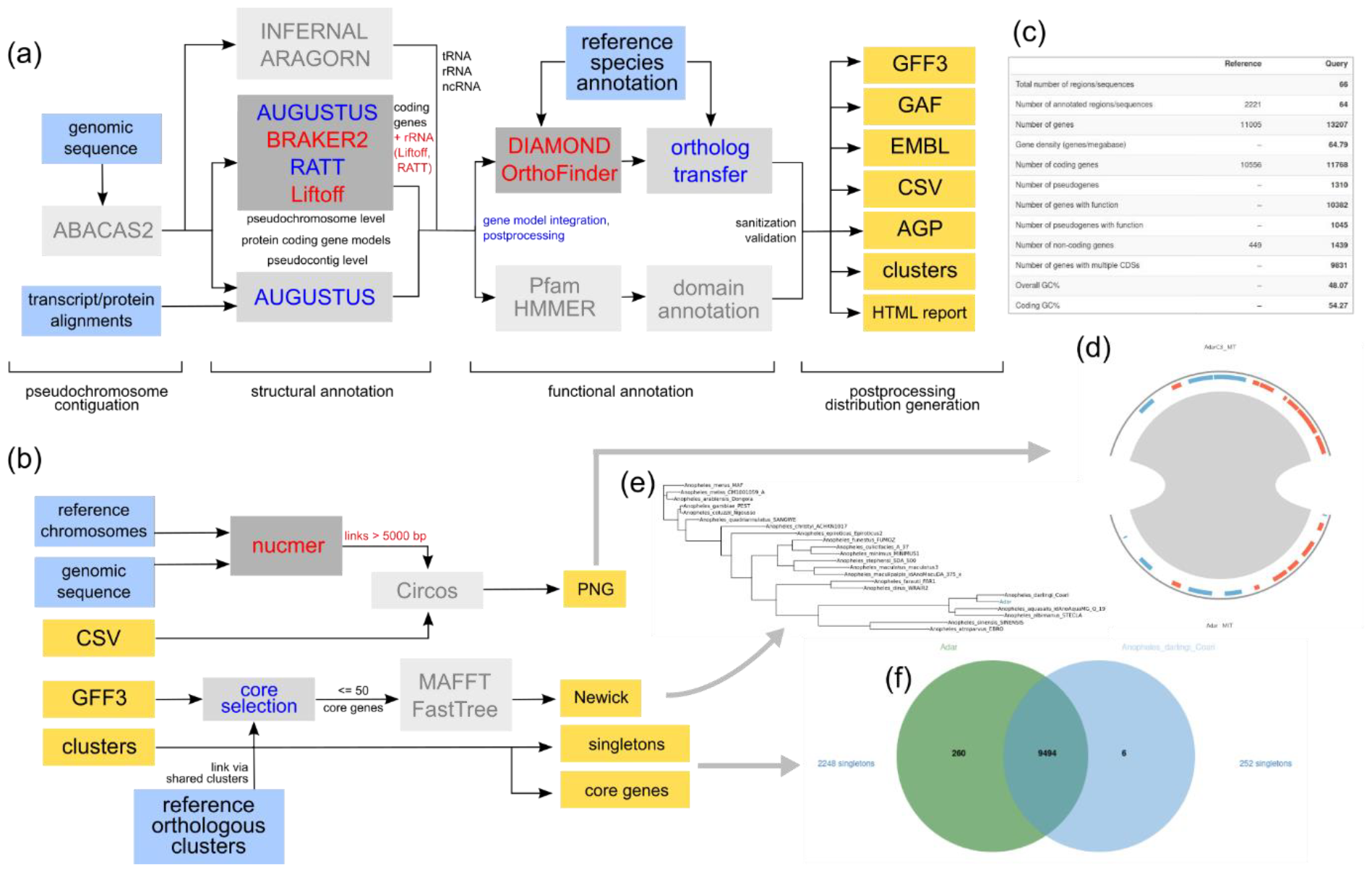
Companion workflows with new components in red and updated components in blue. **(a)** – genome annotation workflow. **(b)** – downstream analysis and visualization workflow. Input files are represented as blue boxes, output files as yellow boxes. All output files are used to construct the result set presented in the web interface, example given of *Anopheles darlingi* assembly target: **(c)** – annotation summary statistics between reference and target; **(d)** – target-reference synteny for mitochondrial chromosome; **(e)** – phylogeny tree placing newly annotated species (here ‘Adar’) in the context of the *Anopheles* genus; **(f)** – interactive Venn diagram summarizing core and species-specific clusters. Summaries of all workflow tools are available in a glossary in the Supplementary Data.

BRAKER2 v2.1.6 (Brůna, et al., 2021) was included in the pipeline and can be used as default option (over AUGUSTUS). Due to the challenges of incorporating a multi-faceted pipeline like BRAKER2, Companion has additional infrastructure to ensure AUGUSTUS is utilised as a backup in the event of process failure. BRAKER is called in protein data mode (see workflow description https://github.com/Gaius-Augustus/BRAKER#braker-with-protein-data), using annotated protein sequences gathered from the reference species family.

Liftoff v1.6.3 (Shumate and Salzberg, 2021) is included as a faster alternative to RATT by default. Default settings are used.

OrthoFinder v2.5.4 (Emms and Kelly, 2019) has replaced OrthoMCL for determining orthologs. It uses DIAMOND (Buchfink, et al., 2021) as a much faster alternative to BLASTP.

MUMmer4.x (Marçais, et al., 2018) v4.0.0rc1 (latest) has enabled replacing BLASTN with nucmer.

For further information and justification for pipeline component upgrades, see Supplementary Data.

### Web server

The web application is implemented in Ruby on Rails with a MySQL database, hosted on a server with 32 CPU cores, 64 GB RAM, as well as over 1 TB storage. It can be accessed at https://companion.ac.uk/. 3 jobs can be run concurrently. Active development is carried out on a development server with matching resources, where full testing of updated features can be carried out without impacting production queues. There is also a build server for integration testing of new Companion release candidates, which has 8 CPU cores, 8 GB RAM and 200 GB storage, with only single job concurrency

For further information about the web server enhancements, see Supplementary Data.

### Testing against alternative web server

To test comparisons with other tools, we used the Companion web server running v2.2.0 and GenSAS v6.0. The input nucleotide sequence fasta file used for each test was identical for both tools. If not otherwise stated, it can be assumed that default settings were used for each test.

For GenSAS, we created an account and selected the closest available AUGUSTUS reference when available for structural annotation. During a naïve first attempt, we selected additional options in preceding tabs (including repeat masking and alignment) on the understanding that these should be expected as part of the pipeline. However, subsequent testing revealed that they dramatically increased runtime with negligible (if any) improvement to accuracy, and so were omitted from final tests.

References for Companion were selected based on their close similarity to the target strain. The target and reference species, along with their VEuPathDB release numbers, can be seen in Supplementary Table S1. Canonical annotations in GFF format were also available to use from each of these releases.

Comparison of outputs with the canonical annotations were performed using GffCompare (Pertea and Pertea, 2020) where such a reference was available, utilising standard metrics such as precision and sensitivity, similar to those used in Holt and Yandell (2011). We focus on four metrics: Nucleotide and exon accuracy, which refer to the proportion of overlap (individual nucleotide bases or exon boundaries, respectively) between a gene prediction and its reference; matching loci, where clusters of exon-overlapping transcripts built for both prediction and reference share at least one match; and total genes (including pseudogenes), which can be used to determine if genes are being overpredicted by either tool.

## RESULTS

### Updates in numbers

Before the improvements to the server, we had around 100 unique users annotating ∼1000 genomes per year. Since Companion version 2 the yearly numbers doubled (annotating ∼2000 genomes). It can be observed that users annotate several genomes or might rerun some annotations with different settings.

### Performance versus GenSAS

To assess the claim that the latest Companion release provides the most user-friendly and accurate annotation platform for a user with minimal additional evidence and a target species from a broad phylogenetic tree, we devised a series of tests to be performed against the web application GenSAS.

First, we tested how easy it is to run both servers. It can be noted that Companion needed just 8 clicks, while GenSAS needed 41 (Supplementary Figure S2) to submit a basic job. Further, in Companion all settings are on one page (Supplementary Figure S2), rather than clicking through different tabs as in GenSAS, making Companion in our view easier to use for users unfamiliar with annotation pipelines.

We ran a series of jobs with matching input genomes on both Companion and GenSAS, aiming to compare annotation outputs to a canonical annotation obtained from VEuPathDB. The assumption in each case was that the user only has the assembly sequence available with no additional evidence, and that a naïve user would tend to choose default settings; a quality annotation should be expected regardless. The input sequences were chosen from Protozoan parasite, Fungi and Vector (see Supplementary Data) species, to highlight Companion’s consistent performance over such phylogenetic breadth of origin.

#### Parasite

##### Plasmodium

As the original release of Companion specialised in annotation of Apicomplexa, including *Plasmodium*, it seemed worthwhile to consider a *Plasmodium* test between strains of close phylogeny, to show both the continued supremacy of Companion for such genomes but also how much it has improved.

*Plasmodium falciparum* Dd2 was chosen as a target due to having a manually curated canonical annotation available (Otto, et al., 2018), and its close phylogeny to the also-well-annotated and established reference *Plasmodium falciparum* 3D7 (Bohme, et al., 2019). This represents a comparison of very similar genomes with divergent subtelomeric regions (5%). AUGUSTUS as implemented in GenSAS doesn’t contain a *Plasmodium* training set, so we were forced to use GeneMarkES for (*ab initio*) structural annotation – this was the only structural tool that didn’t require additional evidence.

Overall, Companion outperforms GenSAS, see Table 1. Especially in the matching loci, the advantages for the reference transfer can be seen. Also, GenSAS fails to find >10% of the genes which is due to the self-training GeneMarkES. Supplementary Figure S3 shows that most gene orthogroups are found by Companion, with many missed by GenSAS. The lower number of genes and low matching loci, however high nucleotide accuracy, in GenSAS can be partially explained by its incorrect merging of several transcripts as a single gene, see Supplementary Figure S4. In the same figure it can also be seen how GenSAS fails to call smaller exons.

#### Fungi

##### Candida

Companion’s reference database has been expanded to include all available species from FungiDB, which includes fungal pathogens of humans, animals and plants, as well as model organisms. (Supplementary Table S1). We decided to use *Candida dubliniensis* CD36 (Jackson, et al., 2009) as a target species against the reference *Candida albicans* SC5314 (Jones, et al., 2004) as this would allow testing performance where there is slightly more phylogenetic distance (albeit the same genus). Once more, there was a well-annotated canonical annotation hosted on FungiDB.

This time, we noted that the AUGUSTUS training set offered by GenSAS contains a *Candida albicans* reference, so this was selected for structural annotation. The results of an almost identical run (but with RATT instead of Liftoff and using Companion v2.0.10) can also be seen with all outputs and visualisations at https://companion.gla.ac.uk/examples. Some of these outputs are summarized in Supplementary Figure S6, showing how closely related the genomes are and how well the interface helps the user to understand the differences between the reference and query.

As GenSAS has a pretrained model, the difference between the two runs is not as striking, e.g. Companion is just 5% better in the matching loci (Table 1). Interestingly, for GenSAS the number of genes is low (∼600 less than the reference), however the nucleotide accuracy is higher than in Companion. This can be explained by GenSAS merging genes (see Supplementary Figure S4). Once again, consistent improvement in the standardised metrics can be observed for Companion.

#### Vector

##### Anopheles

The final comparison was with the *Anopheles darlingi* vector (Table 1 and Supplemental Data). In terms of comparison between Companion and GenSAS, the picture was similar to the Fungi test. However, both methods overpredicted the number of genes and it can be seen that the overall performance dropped. As a new metric we used Pfam domains (Supplementary Figure S7), where we could show that the genes predicted by Companion have a higher coverage.

In conclusion, in all tests Companion outperforms its competitor over all criteria. Using the default settings, Companion is also faster on the web server versus the full GenSAS pipeline (Table 1) and is easier to use.

## DISCUSSION

Projects associated with the Earth BioGenome Project (EBGP) promise to build the genomes of 4 million species – but how will they be annotated? The Darwin Tree of Life (Sanger and EBI) have their own bespoke pipeline. However, other projects such as the European Reference Genome Atlas (ERGA) (Formenti, et al., 2022) are more community driven with more than 700 groups contributing de novo assemblies. It would be a waste of time, effort and energy for each group to generate their own annotation pipeline. In terms of sustainability, we argue that it is far more economical to have a tested service rather than building and testing new pipelines on an ad-hoc basis. Further, there is a huge bottleneck to upload genomes into the main databases. Although EBI is trying to overcome these issues, as the GFF format is an open file format, including it into databases is a time-consuming and manual process. The consequences are that just genomes without annotation are submitted to the databases, with the risk that the annotation might not be assessable under FAIR principles.

With Companion we present a solution to this dilemma; a service easy to run, freely available, including visualisation options and producing high quality annotations. Although the main strength of Companion comes from the reference-based annotation (Table 1), with the introduction of BRAKER2, we have implemented a tool that can efficiently self-train with protein evidence. However, many groups in the community are interested in the study of diversity of species complexes, understanding the evolution of species, and exploration of specific traits such as antimicrobial resistance. Looking at the assembly statistics, many isolates from a well-annotated reference e.g. *Plasmodium*/*Trypanosome* were annotated in the last years with Companion. Indeed, in the last 6 months alone there have been 275 successful Companion web server jobs with a *Plasmodium* reference. This comes down to goals of researchers to frequently sequence multiple isolates from a single species or from related species, and here the transfer of annotation from a reference is of huge advantage – a unique hallmark of Companion. Although Companion has a predefined reference set, 300 unique users have downloaded the stand alone container since the release of version 2, which allows the use of bespoke reference datasets not within VEuPathDB.

It should be noted that annotation is an open problem! For example, even using stand-alone BRAKER2 or MAKER2 with protein and RNA-Seq evidence, the exon accuracy is generally under 80% for species such as vectors. This explains why for key species such as mouse, human, *Plasmodium*, there are efforts of manual curation. In the absence of those, Companion generates good first pass annotation as shown for the vector annotation (Table 1).

In conclusion, we present an annotation web server for the community to overcome the on-going difficulties of genome annotation. In recent years, we have tripled our usage (Supplementary Data) and expect with the increase of phyla to reach even more communities.

## DATA AVAILABILITY

The Companion web server is freely available to all users at https://companion.ac.uk. The source code for the Companion pipeline can be found at https://github.com/sii-companion/companion. Docker images containing versioned releases of Companion for local containerised running are hosted at https://hub.docker.com/repository/docker/uofgiii/companion.

## ACKNOWLEDGEMENT

We would like to thank Scott Arkison for maintaining the computational infrastructure.

