## Supplementary Data for "Annotation and visualisation of parasite, fungi and arthropod genomes with Companion"

### Companion workflow notes

In this update paper of Companion we aimed to cater for larger genomics (up to 1Gb) and since 2016 several new software tools were published, we updated some components of the pipeline.

In addition to the original AUGUSTUS (Stanke, et al., 2008) and RATT (Otto, et al., 2011), users are now offered BRAKER2 (Brůna, et al., 2021) and Liftoff (Shumate and Salzberg, 2021) as default alternatives, respectively (Figure 1).

BRAKER2 is an enhancement of AUGUSTUS with additional support for gene prediction, with consistently improved outcomes at the expense of longer run times. Companion's reference dataset must now include annotated proteins which are used as input for BRAKER2; these are gathered and formatted automatically by the reference update pipeline (see following section). BRAKER2 then trains GeneMark-EP+ and uses pre-trained AUGUSTUS models to predict genes. Additional scripts have been incorporated into the Nextflow pipeline to ensure any failure during the running / output-parsing of BRAKER2 is accounted for; base AUGUSTUS is used as a backup in these circumstances. This provides additional stability for what is an inherently complex and multi-faceted addition to the Companion workflow.

Liftoff is a recently released faster and more scalable alternative to RATT which "lifts over" reference annotations from a reference sequence aligned to a target sequence using Minimap2 (Li, 2018). The demand for a RATT alternative came about when encountering memory scaling issues when running Companion with some of the larger Vector genome assemblies. However, there is still an advantage in using RATT for certain genomes of greater phylogenetic distance to their references, where we still see improved accuracy. There are additional post-processing scripts for Liftoff and RATT to transfer reference rRNA features as a complement to the pre-existing Infernal prediction.

Functional annotation has also seen substantial changes. OrthoFinder (Emms and Kelly, 2019) is a recent alternative to OrthoMCL for determining orthologues between two sets of proteins. Where OrthoMCL used BLASTP for protein alignment, OrthoFinder uses DIAMOND (Buchfink, et al., 2021) which matches BLASTP for sensitivity but is substantially faster. OrthoFinder also bypasses the OrthoMCL requirement to store results in an SQL database, simplifying the process.

Additional upgrades leading to faster run times include MUMmer4/nucmer (Marçais, et al., 2018) instead of BLASTN for nucleotide alignment when generating Circos plots.

### Glossary of tools

The following is an alphabetical list of all major tools that constitute the Companion pipeline, with a brief description of their function. Their place in the pipeline can be seen in Figure 1.

*ABACAS2* (Assefa, et al., 2009) – ordering and orientating nucleotide sequences along a reference

*ARAGORN* (Laslett and Canback, 2004) – detection of tRNA in nucleotide sequences

*AUGUSTUS* (Stanke, et al., 2008) – gene prediction, either *ab initio* or using extrinsic hints

*BRAKER2* (Brůna, et al., 2021) – gene prediction pipeline, incorporating AUGUSTUS and, another gene prediction tool GeneMark-EP+, trained with protein homology evidence

*Circos* (Krzywinski, et al., 2009) – visualisation tool used for chromosome-level synteny plots

*DIAMOND* (Buchfink, et al., 2021) – protein sequence aligner

*FastTree* (Price, et al., 2009) – phylogenetic tree inference from alignments. Generates Newick file, which is rendered using Phylocanvas

*HMMER* (Eddy, 2011) – detection of sequence homologs using probabilistic hidden Markov models (HMMs).

*INFERNAL* (Nawrocki and Eddy, 2013) – inference of RNA features in nucleotide sequence, used for predicting ncRNA

*Liftoff* (Shumate and Salzberg, 2021) – lifting over of gene features from reference to target sequence

*MAFFT* (Katoh, et al., 2002) – multiple sequence aligner, used in conjunction with FastTree

*Nucmer* (Marçais, et al., 2018) – nucleotide sequence alignment program. Part of MUMmer

*OrthoFinder* (Emms and Kelly, 2019) – detection of gene orthogroups using protein sequences of multiple species

*Pfam* (Mistry, et al., 2020) – database of protein families used by HMMER

*RATT* (Otto, et al., 2011) – transfer of gene models with high synteny from reference to target sequence

### Web server notes

Although a Docker container is available for local operation, and allows users to use their preferred reference genomes, most users continue to interact with Companion via a web interface. Approximately 300 pulls have been reported by DockerHub in the two years since the repository was created, while there has been an order of magnitude greater number of jobs submitted via the web interface in the same time span (>1,000 in just the last 6 months). This has motivated the continued development of the web server infrastructure and code base to ensure a better user experience.

| Source | 2016 | 2019 | 2023 |
|---|---|---|---|
| **amoebadb.org** | * | 7 | 10 |
| **cryptodb.org** | * | 10 | 14 |
| **fungidb.org** | 0 | 112 | 224 |
| **hostdb.org** | 0 | 0 | 9 |
| **microsporidiadb.org** | * | 23 | 39 |
| **piroplasmadb.org** | * | 10 | 13 |
| **plasmodb.org** | * | 23 | 25 |
| **toxodb.org** | * | 14 | 15 |
| **tritrypdb.org** | * | 30 | 37 |
| **vectorbase.org** | 0 | 0 | 52 |
| **TOTAL** | ***62*** | ***229*** | ***438*** |

*Table S1 Number of reference organisms available on Companion web server by source*
** Breakdown unknown; only TOTAL value was quoted in Steinbiss, et al. (2016)*

Companion currently has 438 available references, a 7-fold increase versus the original web server implementation (see Table S1). An automated reference update pipeline has been developed (also using Nextflow DSL1) to accommodate such a large number of references, automatically extracting, formatting and training models for a given reference domain. Source code is available at https://github.com/sii-companion/reference-update. Running the script *bin/run_all.sh* on a dedicated data server, reference species files from every VEuPathDB.org domain are automatically downloaded using the web service interface https://github.com/sii-companion/eupathws, and all files necessary for Companion packaged into a directory. Annotated proteins are gathered from every reference organism of a given VEuPathDB site, combined with the relevant OrthoDB v11 clade (Kuznetsov, et al., 2022), to provide a comprehensive pool of protein evidence as input for BRAKER2. The entire reference dataset is then secure copied to the various Companion servers and loaded into each Rails database using a rake task, which parses unique gene IDs and build numbers to ensure only newly updated/released data is extracted (and so avoid duplication of work). This whole process is carried out twice a year. All references available for the user can be viewed, together with metadata, in a table at https://companion.gla.ac.uk/references/, and individual reference metadata is available as part of the output result files for any completed job.

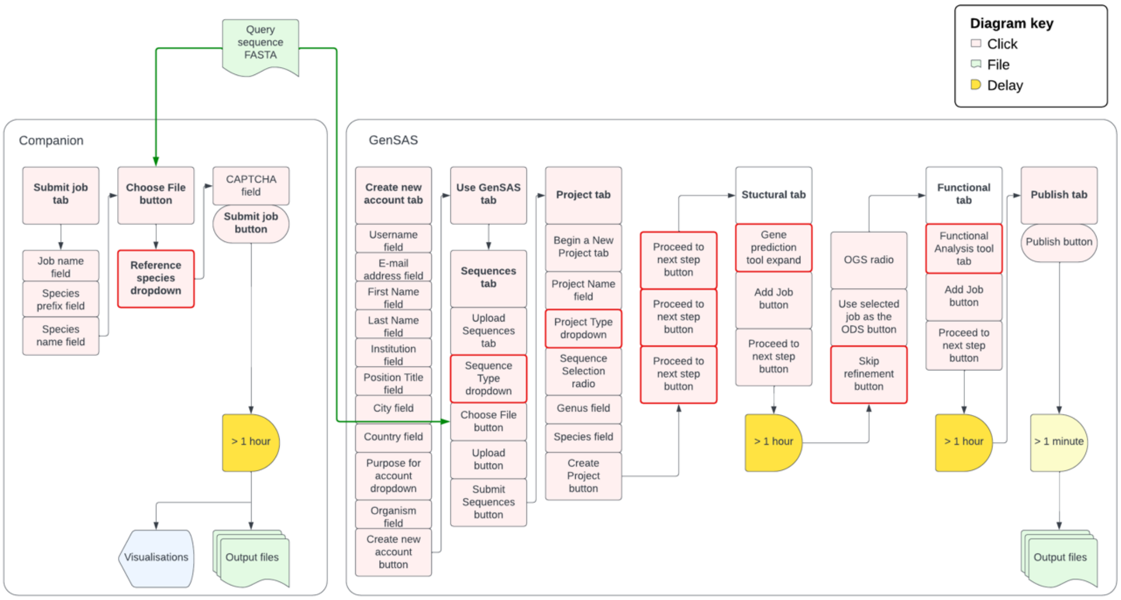


*Figure S1 Minimum clicks required for new user of Companion and GenSAS to perform a basic job with default settings that ensure both structural and functional annotations: 8 and 41, respectively. Clicks highlighted in red require expert insight about the target genome.*

While the core architecture of the Companion web interface still uses the same Ruby on Rails backend as in the first implementation, there have been significant developments to improve job concurrency. A MySQL server is now used instead of SQLite to prevent file locking errors with simultaneous write commands, thus removing the primary barrier to job concurrency on a single server. The effect of this has enabled the use of a single entry point for the Companion web interface housing all available references, where before several URLs were used based on groupings of similar references. 3 jobs can be run concurrently with ease. To allow efficient processing, our production server has 32 CPU cores, 64 GB RAM, as well as over 1 TB of disc space.

Taken together, the core Companion and web server infrastructure changes ensure job completion rates of >95%.

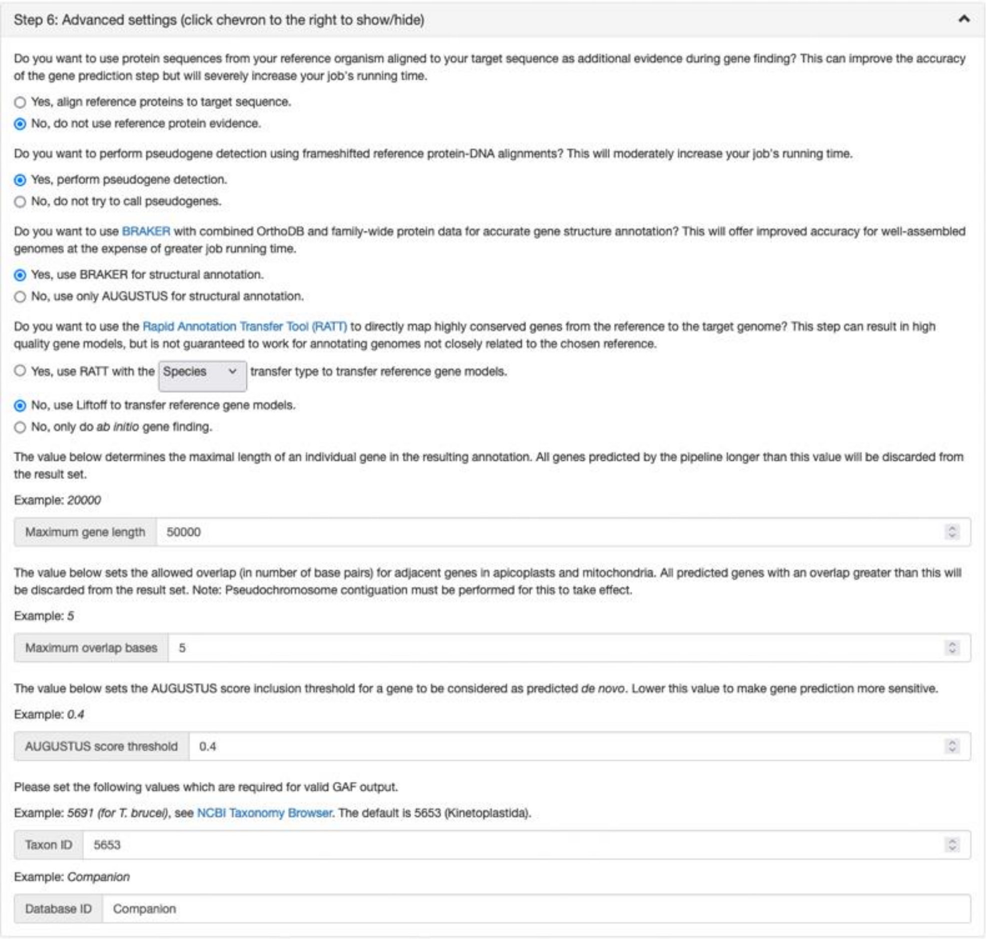


*Figure S2 Companion web server advanced settings for optional refinement.*

### Results notes

| Target | Source | Reference | Source |
|---|---|---|---|
| ***Plasmodium falciparum* Dd2** | PlasmoDB release 65 | *Plasmodium falciparum* 3D7 | PlasmoDB release 9 |
| ***Candida dubliniensis* CD36** | FungiDB release 60 | *Candida albicans* SC5314 | FungiDB release 60 |
| ***Anopheles darlingi* assembly idAnoDarlMG-H_01** | VectorBase release 65 | *Anopheles darlingi* Coari | VectorBase release 49 |

*Table S2 Data sources for target and reference of three comparisons.*

#### Parasite: *Plasmodium*

Companion completed in ~2.5 hours. Selecting options from most of the pre-structural tabs (like repeat masking with EvidenceModeller and alignment with BLASTn), GenSAS took 6h. Excluding the mentioned settings, GenSAS completed in ~1 hour, (although the higher runtime in Companion was mostly accounted for by ncRNA prediction with Infernal – we neglected to choose additional ncRNA annotation tools as part of the GenSAS job).

To be able to make the *Plasmodium falciparum* Dd2 comparison, we excluded UTR features that were transferred by Liftoff from the reference, as the current annotation did not have them annotated.

Overall Companion is very effective in annotating the *Plasmodium* genome however there are some over-predictions. Looking into these, they appear to mostly be in the teleomeric regions where repeats are looking like genes. Although these genes are easy to detect in tools like Artemis (Carver, et al., 2012), it would still require manual intervention to obtain a perfect annotation.

Companion was run with default settings (BRAKER2 and Liftoff) and in a second run with RATT (Strain setting), owing to similarity of target and reference (see Table S).

| Organism | Metric | Companion (Liftoff) | Companion (RATT) |
|---|---|---|---|
| *Plasmodium falciparum* (5,642 total genes) | Nucleotide accuracy (%) | 99.35 | 99.4 |
| | Exon accuracy (%) | 95.05 | 94.9 |
| | Matching loci (%) | 96.22 | 96.02 |
| | Total genes predicted | 5,800 | 5,791 |
| *Candida albicans* (6,459 total genes) | Nucleotide accuracy (%) | 98.35 | 98.65 |
| | Exon accuracy (%) | 89.50 | 90.15 |
| | Matching loci (%) | 95.54 | 96.03 |
| | Total genes predicted | 6,206 | 5,833 |

*Table S3 Comparable results when using RATT for gene finding rather than default Liftoff.*

Performing the same job on legacy Companion (version 1.0.2) (although naturally with AUGUSTUS/RATT rather than BRAKER2/Liftoff) showed similarly high accuracy to the current annotation but took almost four times longer to run (~9 hours). As the number of processors was increased from 8 to 32, we can see a near linear decrease in runtime due to parallelisation. Comparisons to the current annotation showed similarly high sensitivity between both runs, but Companion version 2 matched 35 more loci (0.6%). There have been notable improvements in capturing apicoplast and mitochondrial ncRNA in the latest Companion compared to version 1. It captured 4 apicoplast rRNA and 21 mitochondrial rRNA, all of which were omitted in the legacy run of Companion as well as GenSAS (using additional RNAmmer tool in pipeline), see Figure S5.

Overall, the new version of Companion not only outperforms existing tools not built for parasite genomics, but also shows improvement in speed and accuracy to its previous version.

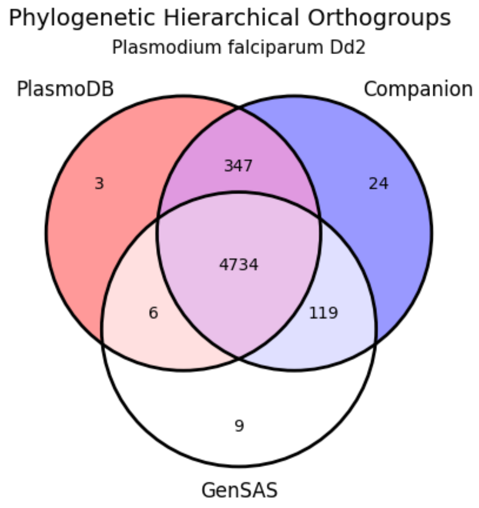


*Figure S3 Venn diagram of shared Hierarchical Orthogroups found by OrthoFinder. Notice greater number of orthogroups shared by Companion and current annotation (PlasmoDB) 347, versus only 6 shared exclusively between GenSAS and PlasmoDB. Note also 119 orthogroups suggesting novel genes.*

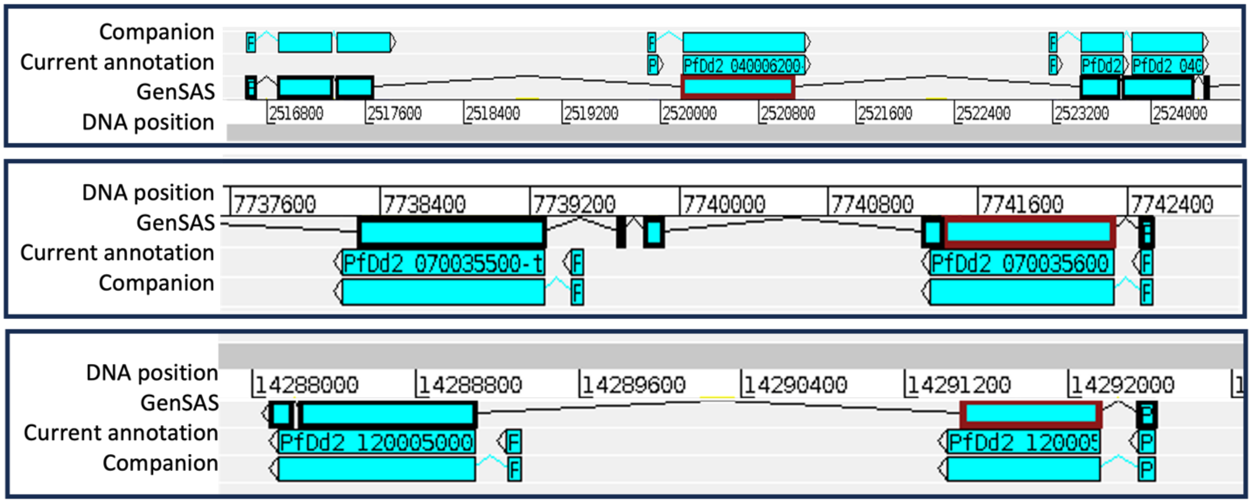


*Figure S4 GenSAS (inner track) frequently incorrectly merges genes as one (see highlighted CDS with intron bridging)*. It also misses smaller exons. Interestingly, in the first panel, the current annotation has a missing gene.

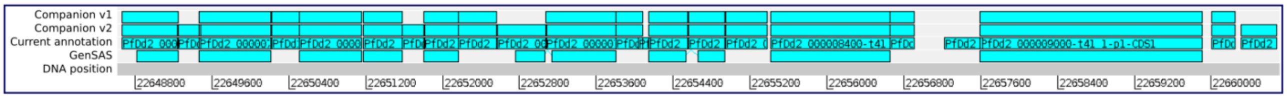


*Figure S5 Several apicoplast genes matched by Companion v2 and current annotation (middle two tracks) but missed by GenSAS (bottom track) and Companion v1. The two genes missed by Companion v2, PfDd2_000008050 and PfDd2_000008950, were due to an overlap with an adjacent gene of 34 and 16 bases, respectively; outside the default value of Companion's new maximum overlap setting (see Figure S2).*

### Fungi: *Candida*

Both Companion and GenSAS jobs completed in approximately 1 hour. Like before, additional pipeline tools in GenSAS (such as RepeatModeller, BLASTn alignment) resulted in much greater run times for modestly worse performance overall. The output results of Companion can be seen in Figure S6.

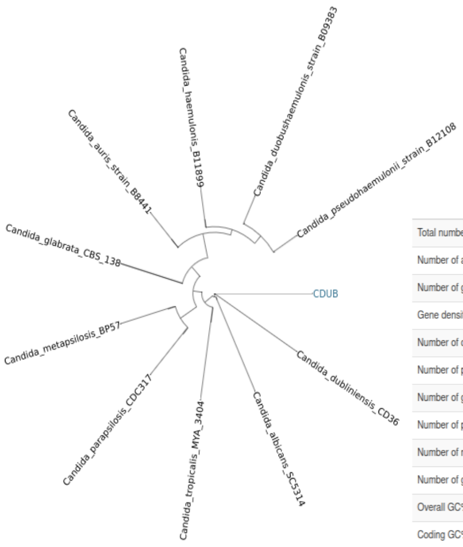


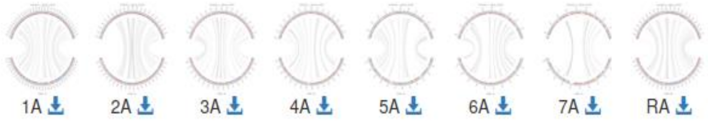


| | Reference | Query |
|---|---|---|
| Total number of regions/sequences | 9 | 8 |
| Number of annotated regions/sequences | 9 | |
| Number of genes | 6459 | 5930 |
| Gene density (genes/megabase) | – | 399.02 |
| Number of coding genes | 6212 | 5833 |
| Number of pseudogenes | – | 193 |
| Number of genes with function | – | 5734 |
| Number of pseudogenes with function | – | 190 |
| Number of non-coding genes | 247 | 97 |
| Number of genes with multiple CDSs | – | 448 |
| Overall GC% | – | 33.25 |
| Coding GC% | – | 34.89 |

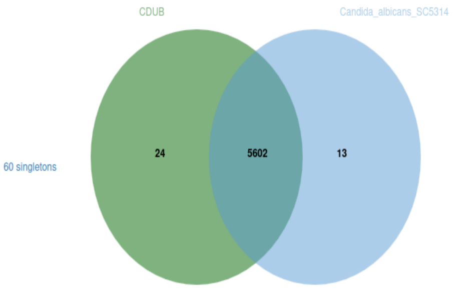


*Figure S6 Visualisations of fungi example run, showing radial phylogenetic tree, synteny, orthology and statistics.*

### Vector: *Anopheles*

Finally, to highlight the expansion of Companion to allow larger vector genomes, we decided to annotate an unpublished *Anopheles darlingi* assembly (idAnoDarlMG-H_01, accession PRJEB53253) against the same-species reference *Anopheles darlingi* Coari (Marinotti, et al., 2013).

Companion was once again run with BRAKER2 for structural annotation. This time, Liftoff was an obvious choice over RATT to account for the better memory scaling of Liftoff for larger genomes and its impressive performance lifting over genes from target and reference of high similarity. The GenSAS AUGUSTUS training set only includes two vector species: *Aedes aegypti* and *Drosophila melanogaster*, so we were forced once more to use GeneMarkES (ab initio) for structural annotation. The use of GeneMarkES ensured that the GenSAS job completed in ~3 hours versus the Companion job's ~9 hours runtime.

Mitochondrial gene transfer was also carried out by Companion. All 13 orthologous mRNA were transferred from the reference AdarC3_MT sequence, and an additional 2 tRNA were predicted. None of these non-coding RNA are available in the current annotation, nor were they predicted by GenSAS.

Table 1 metrics show generally favourable results for Companion versus GenSAS, although neither tool achieves comparably high outcomes to the previous tests, implying there's still work to be done in improving Vector annotation across the board. It is important to note that quality annotation of vector genomes is relatively few and far between (the study for the chosen assembly remains unpublished), and so the notion of such a reference annotation being a valid "truth set" comparison is questionable. This motivated running GFFCompare after filtering for only CDS features in both the reference and query annotations, where the inclusion of UTRs in the current annotation appeared to seriously hamper the metrics for both tools.

A method of validation that doesn't require reference annotation is BUSCO completeness (Seppey, et al., 2019), which assesses protein sequences against a database of proteins from a relevant lineage. Using lineage database insecta_odb10, proteins output by Companion achieved 95.4% complete (C), 0.7% fragmented (F) and 3.9% missing (M). This compares favourably to the GenSAS output which achieved 94.1% C, 1.1% F and 4.8% M. The more specific lineage database diptera_odb10 saw scores of 93.9% C, 1.4% F and 4.7% M for Companion, versus comparable results of 93.2% C, 1.7% F and 5.1% M for GenSAS. The current annotation idAnoDarlMG-H_01 assembly proteins (from VectorBase) achieved 99.1% C, 0.1% F and 0.8% M against diptera_odb10, for comparison.

Another alternative method of validation that also considers proteins is coverage of genes that contain a Pfam domain annotation (this was used successfully in Holt and Yandell (2011)). The results, including a breakdown of some of the more prominent functions, can be seen in Figure S7. There is a modest improvement in Companion of 2.2% against GenSAS. The addition of InterProScan to perform protein matching of Pfam domains added ~10.5 hours to the overall GenSAS runtime. HMMER v3.3 (the same as used in the Companion pipeline) was run independently on the current annotation proteins to determine their Pfam coverage.

It should be noted that we tested various approaches on the command line to improve the annotation, using different input of RNA-Seq data, training with proteins of the Vector domain only. However, Companion always returned the best annotation results.

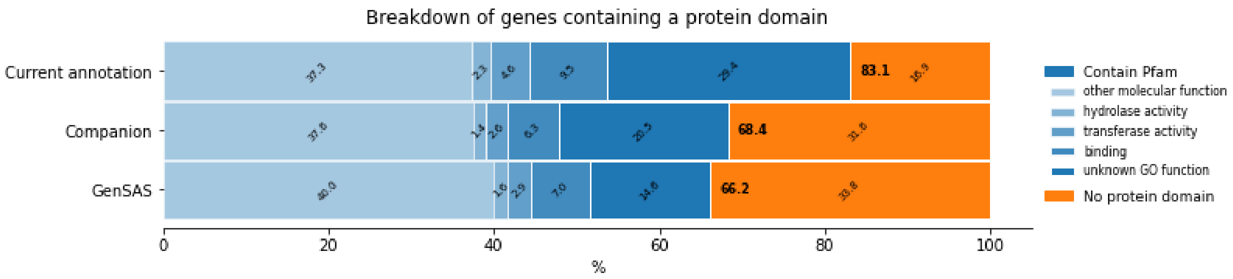


*Figure S7 Percentage of genes that were annotated with a Pfam domain.*
